# A brief report of sleep and circadian rhythm quotas in a population of dog owners in North Carolina, USA

**DOI:** 10.1101/2021.01.21.427658

**Authors:** Ujas A. Patel, Margaret E. Gruen, David R. Samson

**Affiliations:** Department of Anthropology, University of Toronto, Mississauga, ON, CA; Department of Clinical Sciences, College of Veterinary Medicine, North Carolina State University, Raleigh, NC, USA

**Keywords:** sleep, actigraphy, circadian rhythms, dog-owners

## Abstract

**Objectives:** To characterize sleep and circadian rhythms of a sample population of healthy, dog-owning adults from North Carolina, USA.

**Methods:** Actigraphy was used to analyze sleep-wake patterns in forty-two dog owners from the Raleigh area in North Carolina. Sleep quotas, including sleep duration, efficiency, and fragmentation were measured alongside a Non-parametric Circadian Rhythms Analysis (NPCRA) to quantify strength, consistency, and fragmentation of rhythms.

**Results:** Compared to females, males demonstrated later sleep onset and sleep end (*p*<0.01), greater wake after sleep onset and sleep fragmentation (*P*<0.001), and lower sleep efficiency (*p*<0.001). The NPCRA revealed comparable relative amplitude (strength) and interdaily stability (consistency), yet less intra-daily variability (fragmentation), than previously reported post-industrial samples.

**Conclusions:** This study adds to the current data available on sleep and circadian rhythms in discrete human populations and highlights the need for more research characterizing cross-cultural sleep and circadian rhythmicity.

## INTRODUCTION

With advances in biometrics and the widespread use of actigraphy outside of laboratory settings, the ability to measure the breadth of human sleep expression in multiple ecological, technological, and cultural contexts has been the hallmark of sleep research in the 21^st^ century (Carskadon and Dement 2005, Roenneberg 2013, Samson 2020). Despite such advances in research, we have yet to answer many fundamental questions about how human sleep varies cross-culturally and globally.

To address this challenge, work describing sleep and circadian rhythms in multiple populations dwelling in both large and small-scale societies and practicing different modes of economic production (Roenneberg 2013, Samson 2020) should be performed. Current research only contains a handful of studies that characterize sleep in discrete human populations (Roenneberg 2013, Worthman and Melby 2002), and of those, only a few also characterize circadian rhythmicity (Rock, Goodwin et al. 2014, Beale, Pedrazzoli et al. 2017, Samson, Crittenden et al. 2017, Samson, Manus et al. 2017, Smit, Broesch et al. 2019). In fact, despite the overwhelming majority of sleep studies having been conducted in U.S. and European settings, samples from this context are still limited in socio-demographic breadth. Thus, it is crucial to report not only sleep quotas but also measures of circadian rhythmicity to understand the factors that drive sleep-wake regulation in varied human populations.

Parallel studies of sleep in dog owners and dogs is an area of great translational potential; dogs and humans share risks for many conditions that affect sleep. As part of a study establishing a baseline sleep-wake cycle and activity patterns using actigraphy and functional linear modeling (FLM), for healthy, adult companion dogs (Woods, Li et al. 2020), we recorded sleep in a population of working professionals who also owned dogs. Thus, the purpose of this study is to add to the developing database of human sleep in varied contexts by characterizing sleep and circadian rhythms of a sample population of healthy, dog-owning adults from North Carolina, USA.

## METHODS

### Study location and Participants

A sample of dog owners was recruited from the Raleigh area in North Carolina (35.7796° N, 78.6382° W) as a part of a multi-phase project designed to generate baseline sleep-wake cycle and activity patterns for companion dogs. Study participants were recruited from the area surrounding the North Carolina State University College of Veterinary Medicine (NCSU CVM) through email and social media. To determine participant eligibility, participants between the age of 18 and 62 years of age were required to take the Pittsburgh Sleep Quality Index (PSQI) (Buysse, Reynolds et al. 1989). A PSQI score of less than or equal to five was classified as a normal sleeper. Participants were screened as being able to participate if they did not have a medical diagnosis of a sleep disorder and were not classified on the PSQI as having poor sleep. Consent from eligible participants was attained by way of written participation forms, which were approved by way of the Research Ethics Board of Canada. The approval was attained under the RIS Human Protocol Ethics # 35696.

The study was conducted during the winter season between January 23^rd^ and February 17^th^, 2019. Day length during this time of year ranged between 10.22 and 11.00 hours. Sunrise occurred between 06:58 and 07:20, and sunset occurred between 17:33 and 17:58 hours. Day length was recorded from Sunset-Sunrise (https://sunrise-sunset.org/). A final sample of 42 dog owners was included for data collection and subsequent data analysis. The sample population is representative of a typical western, educated, industrialized, rich, and democratic – so called “WEIRD” population (Henrich, Heine et al. 2010). Participants ranged from 21 years to 62 years of age with an average age of 33.5 ± 10.4 years. The majority had regular daytime schedules, working five days of the week with the exception of six individuals (three each worked four and seven days per week).

### Actigraphy

Motionwatch 8 actigraphs (CamNtech) were used for data collection, with all devices configured to generate data in one-minute epochs. Participants were asked to press the event marker on the Motionwatch preceding any sleep events throughout the study duration. Actigraph data were scored using CamNtech MotionWare 1.2.23 software, which uses a sleep detection algorithm to generate sleep quotas based on actigraphic counts. The software also has a nap analysis function, with adjustable parameters, to detect periods of inactivity as a nap. We followed parameter values that have been validated for use in the field environments using actigraphy (Samson, Yetish et al. 2016) to define the criteria to score a period of inactivity as napping (i.e., daytime sleep).

### Statistical Analyses

Statistical analyses were conducted using *R* version 1.3.1093 (2020). Descriptive statistics were generated to characterize sleep in the sample population. Male and female sleep quotas were presented and compared using a Wilcoxon Rank Sum test. To characterize circadian rhythms in the sample population, Nonparametric Circadian Rhythm Analysis (NPCRA) (Van Someren, Swaab et al. 1999) was performed on the data generated from the study sample.

## RESULTS

Sleep quotas characterizing the North Carolina (NC) sample were calculated for each sex (Table 1). A sum of 472 days of day and night activity data were generated. A total of 42 participants were included in the study with 27 females (mean age: 32.6 ± 10.0 years) and 15 males (mean age: 35.3 ± 10.2 years). Males were characterized by later sleep onset and sleep end (*p* < 0.01), greater wake after sleep onset (WASO) and sleep fragmentation (*p* < 0.001), and lower sleep efficiency (*p* < 0.001) (Table 1). Results for the NPCRA are presented in Table 2. The mean NPCRA for the sample (*n* = 42) was 26.4 days. The analysis reveals a relative amplitude (RA) of 0.90 ± 0.05, demonstrating high circadian amplitude in the NC sample. Parameters measuring the consistency and fragmentation of circadian rhythms in the study population reveal an interdaily stability (IS) of 0.46 ± 0.11 and intra-daily variability (IV) of 0.57 ± 0.21, respectively.

**Table 1.** Study sample characteristics for full sample by sex (male n = 15, female n = 27). Data are presented as mean (standard deviation)

| Sleep-wake measure | Men | Women | Total | Significance |
| --- | --- | --- | --- | --- |
| Sleep onset (hh:mm) | 23:23 (1:41) | 22:58 (1:23) | 23:07 (1:30) | $W = 36756$;<br>$p < 0.01$ |
| Sleep end (hh:mm) | 07:25 (1:37) | 07:05 (1:36) | 07:12 (1:37) | $W = 37268$;<br>$p < 0.01$ |
| Sleep latency (hour) | 0.25 (0.33) | 0.21 (0.24) | 0.23 (0.27) | $W = 43138$;<br>$p = 0.8114$ |
| Time in bed (hour) | 8.36 (1.57) | 8.41 (1.44) | 8.39 (1.49) | $W = 43888$;<br>$p = 0.5497$ |
| Sleep duration (hour) | 8.02 (1.56) | 8.12 (1.38) | 8.08 (1.45) | $W = 44490$;<br>$p = 0.3749$ |
| Wake after sleep onset (hour) | 1.49 (0.60) | 1.21 (0.54) | 1.31 (0.58) | $W = 30478$;<br>$p < 0.001$ |
| Sleep efficiency (%) | 78.44 (6.54) | 82.39 (5.90) | 80.98 (6.42) | $W = 57492$;<br>$p < 0.001$ |
| Sleep fragmentation | 31.16 (10.15) | 26.81 (11.04) | 28.36 (10.92) | $W = 32084$;<br>$p < 0.001$ |
| Cumulative night-time activity | 99484 (30659) | 85828 (30265) | 90705 (32691) | $W = 153$;<br>$p = 0.2009$ |
| Daily nap ratio | 0.28 (0.21) | 0.32 (0.25) | 0.30 (0.23) | $W = 209$;<br>$p = 0.8745$ |

**Table 2.** Nonparametric Circadian Rhythm Analysis (NPCRA)

| Parameter | Mean (SD) | Definitions |
| --- | --- | --- |
| Interdaily stability (IS) | 0.46 (0.11) | Degree of resemblance between the activity patterns on individual days; ranges from 0 to 1 and may typically be about 0.6 for healthy adults |
| Intra-daily variability (IV) | 0.57 (0.21) | Quantifies the fragmentation of periods of rest and activity; ranges from 0 to 2 and typically is $<1$ for healthy adults, with higher values indicating a more fragmented rhythm |
| L5 | 747.05 (359.65) | All days of data are overlaid and averaged in 24-h periods. The L5 average provides the mean activity level for the sequence of the least five active hours, indicating how inactive and regular sleep periods are. |
| M10 | 15368.24<br>(3599.95) | All days of data are overlaid and averaged in 24-h periods. The M10 average provides the mean activity level for the sequence of the most active hours, indicating how active and regular the wake periods are. |
| Relative amplitude (RA) | 0.90 (0.05) | Calculated by dividing amplitude by the sum of L5 and M10; ranges from 0 to 1, with higher values indicating higher amplitude of the rhythm |
| L5onset | 12:27 (1:09) | Onset time of the five most restful consecutive hours |
| M10onset | 08:26 (3:31) | Onset time of the 10 most active consecutive hours |

## DISCUSSION

In this study, we characterized sleep and circadian rhythms in a sample population of NC dog owners. We found significant differences between male and female participants in their sleep onset and end times, duration of WASO, amount of sleep fragmentation, and sleep efficiency. Similar sex differences in WASO have been previously reported in other post-industrial populations (Ohayon, Carskadon et al. 2004, Goel, Kim et al. 2005). By contrast, most sleep studies that include sex differences report mixed observations for all other sleep variables (Ohayon, Carskadon et al. 2004, Mong and Cusmano 2016). The NC sample reinforces previously reported trends in sleep between post-industrial populations and small-scale societies, where the present sample demonstrates a longer sleep duration (Prall, Yetish et al. 2018, Smit, Broesch et al. 2019, Yetish and McGregor 2019), shorter sleep latency (De la Iglesia, Fernández-Duque et al. 2015, Samson, Crittenden et al. 2017, Samson, Manus et al. 2017), and greater sleep efficiency (Samson, Crittenden et al. 2017, Samson, Manus et al. 2017, Prall, Yetish et al. 2018) than small-scale societies. The NPCRA on the sample data (Table 2) reveals comparable RA (strength) and IS (consistency), yet less IV (fragmentation), than previously reported post-industrial samples (Rock, Goodwin et al. 2014). Yet, these circadian measures await future comparative analysis. Future work should incorporate multiple measures of circadian strength into a Circadian Function Index (Ortiz-Tudela, Innominato et al. 2016, Ortiz-Tudela, Martinez-Nicolas et al. 2010) to model the influence of varied sleep ecologies and to better understand the role of societies’ influences on sleep and circadian function.

This study is not without limitations. Polysomnography is considered the “gold-standard” for sleep research – thus the use of actigraphy data to characterize sleep should be used with caution. Wrist actigraphy infers sleep based on the lack of movement (Samson 2020, Martin and Hakim 2011); if an individual is physiologically awake but performs little to no movement, the actigraph scores this as ‘asleep’. Accordingly, actigraphy has a bias towards overestimating certain sleep variables such as naps, total sleep time, sleep onset latency, etc. (Samson 2020, Martin and Hakim 2011). Nonetheless, actigraphical detection of sleep-wake patterns have been validated against polysomnography by several studies with high concordance and >90% sensitivity in healthy individuals (Samson 2020, Martin and Hakim 2011).

In conclusion, new technologies have made the study of sleep more viable, allowing for novel opportunities to study sleep and circadian rhythms in different human populations (Samson 2020, Deboer 2007). Generating data-sets characterizing sleep and circadian rhythmicity in human populations, that differ in ecology, culture, and lifestyle, is essential to understanding the role of sleep in a cross-cultural context – ultimately improving the capacity to test evolutionary hypotheses about the function of human sleep (Beale, Pedrazzoli et al. 2017, Samson, Crittenden et al. 2017, Samson, Manus et al. 2017, Nunn, Samson et al. 2016). Additionally, with the growing number of societies where companion animals live and sleep in the same environmentally buffered domiciles, future work should model the interaction of pet-human sleep and co-sleeping environments. In sum, this study adds to the current data available on sleep and circadian rhythms in discrete human populations. Given the handful of studies that report such data, this study promotes the need for more research characterizing sleep and circadian rhythms in healthy human populations across the globe.

## ACKNOWLEDGEMENTS

We thank the dog owners who participated in the study for their time and commitment. We also thank Andrea Thomson for the collection and management of project data. Finally, we thank the SSHRC Insight Development Grant: 430-2018-00018 for funding support.

